# BRD2 and BRD3 genes independently evolved RNA structures to control unproductive splicing

**DOI:** 10.1101/2023.10.08.561383

**Authors:** Marina Petrova, Sergey Margasyuk, Margarita Vorobeva, Dmitry Skvortsov, Olga Dontsova, Dmitri D. Pervouchine

## Abstract

The mammalian *BRD2* and *BRD3* genes encode structurally related proteins from the bromodomain and extraterminal domain (BET) protein family. The expression of *BRD2* is regulated by unproductive splicing upon inclusion of exon 3b, which is located in the region encoding a bromodomain. Bioinformatic analysis indicated that *BRD2* exon 3b inclusion is controlled by a pair of conserved complementary regions (PCCR) located in the flanking introns. Furthermore, we identified a highly conserved element encoding a cryptic poison exon 5b and a previously unknown PCCR in the intron between exons 5 and 6 of *BRD3*, however outside of the homologous bromodomain. Minigene mutagenesis and blockage of RNA structure by antisense oligonucleotides demonstrated that RNA structure controls the rate of inclusion of poison exons. The patterns of *BRD2* and *BRD3* expression and splicing show downregulation upon inclusion of poison exons, which become skipped in response to transcription elongation slowdown, further confirming a role of PCCRs in unproductive splicing regulation. We conclude that *BRD2* and *BRD3* independently acquired poison exons and RNA structures to dynamically control unproductive splicing. This study describes a convergent evolution of regulatory unproductive splicing mechanisms in these genes providing implications for selective modulation of their expression in therapeutic applications.

## Introduction

The bromodomain and extraterminal domain (BET) protein family, a subgroup of the bromodomain protein superfamily [1, 2, 3], consists of four related chromatin readers (*BRD2*, *BRD3*, *BRD4*, and *BRDT*) with broad specificity on transcriptional activation of immunity-associated genes [4, 5]. They contain two homologous tandem bromodomains, which recognize and bind acetylated lysine residues, an extra-terminal (ET) domain, and two small conserved motifs located between the two bromodomains and between the second bromodomain and the ET domain [6]. Abnormal expression or loss of function in BET proteins is associated with pathological processes, including immune and inflammatory diseases, and is being increasingly considered as a prominent therapeutic target [7].

BET family members have evolved in a series of duplications that occurred before vertebrate radiation [8]. A high degree of sequence similarity, interactions with many of the same transcription factors [9, 10, 11, 12], and overlapping epigenetic binding profiles [13] suggest that BET proteins have similar cellular activities. However, despite these shared characteristics, there appears to be no functional redundancy between them [14]. The fact that BET members cannot compensate for each other is emphasized by the lethal consequences of *BRD2* [15] and *BRD4* [16] deletion. Unlike *BRD3*, *BRD2* doesn’t have reprogramming activity [17] and its depletion doesn’t lead to cell death [13]. The depletion of *BRD2* but not *BRD3* or *BRD4* reverts the sigma-2 receptor up-regulation induced by cholesterol deprivation [18]. Yet, the molecular mechanisms underlying BET-specific functions remain largely unresolved.

The bromodomain proteins, and BET family members in particular, can be regulated at the post-transcriptional level through a mechanism called unproductive splicing, in which the mRNA is triggered to degradation by the nonsense-mediated decay (NMD) as a result of regulated alternative splicing [19]. The inclusion of exon 3b, which is embedded in conserved intronic sequences, introduces a frame shift and thus a premature termination codon (PTC) in the *BRD2* transcript [20, 21]. Such exons are called poison exons because they are normally skipped but trigger mRNA degradation upon inclusion. Another example is the *BRD9* gene, which contains a poison exon that leads to mRNA degradation in SF3B1-mutant tumors [22]. Multiple cross-regulatory unproductive splicing networks tend to evolve among paralogs often containing ultraconserved elements around poison exons [23].

In this work, we questioned whether other BET family members, which are *BRD2* paralogs, could be regulated through unproductive splicing and whether this regulation is evolutionarily conserved. We identified a previously unknown cryptic poison exon in *BRD3*, which however resides in a non-homologous intron with respect to the poison exon in *BRD2*, and predicted bioinformatically that both poison exons are surrounded by pairs of conserved complementary regions (PCCR) capable of forming stable RNA structures. Using blockage of RNA structure by antisense oligonucleotides and minigene mutagenesis, we demonstrated that base pairings between these PCCRs indeed control poison exon inclusion. Furthermore, we characterized unproductive splicing patterns in these genes using panels of healthy human tissue transcriptomes from the Genotype Tissue Expression (GTEx) project and tumor transcriptomes from the The Cancer Genome Atlas (TCGA). Finally, we described characteristic RNA-structure-specific patterns in unproductive splicing response to the transcription elongation slowdown.

## Results

### Phylogenetic analysis

A study by Paillisson *et al.* suggested that vertebrate BET family consists of four separate orthologous groups corresponding to *BRD2*, *BRD3*, *BRD4*, and *BRDT*, respectively, with the highest degree of similarity between *BRD2* and *BRD4* [8]. In repeating this analysis, we constructed a phylogenetic tree of bilaterian homologs of *BRD2* from eggNOG database [24], which confirmed the separation of four orthologous groups, however suggesting the most recent divergence of *BRD2* and *BRD3*, not *BRD2* and *BRD4* (a restricted tree is shown in Figure 1A). Interestingly, zebrafish has evolved two copies of *BRD2*, while mouse maintained only one (Figure S1). The hypothesis that the closest homolog of *BRD2* could be *BRD3* is also evidenced by the relationship between phylogeny and expression profiles of these genes [8], and by the fact that *BRD2* and *BRD3* both have a shorter C-terminal domain as compared to *BRD4* and *BRDT* [25]. The phylogenetic trees derived from multiple sequence alignments of the two bromodomains also confirm the highest similarity of *BRD2* to *BRD3*, not to *BRD4* (Figure S2).

**Figure 1:**
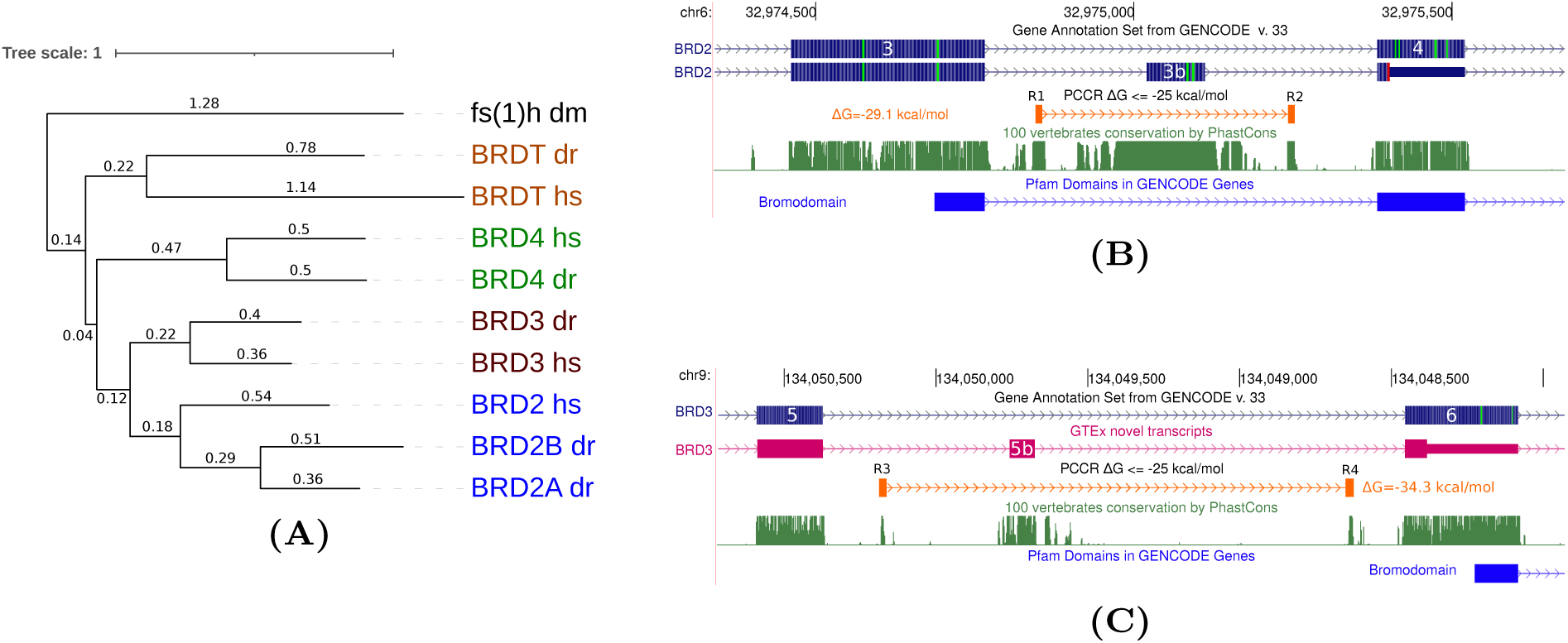
The BET family. **(A)** The phylogenetic tree of bilaterian homologs of *BRD2* from eggNOG database restricted to *H. Sapiens* and *D. rerio*. The *fs(1)h* protein from *D. Melanogaster* was used as an outgroup. The numbers on the branches denote the average number of aminoacid substitutions per site. **(B)** The inclusion of exon 3b in *BRD2* results in a PTC in the downstream exon. The exon is surrounded by a PCCR formed by R1 and R2 with Δ*G*=-29.1 kcal/mol. Exon 3b is located within a region encoding the bromodomain 1. **(C)** Cryptic exon 5b in *BRD3* is unannotated and can be detected in a few human tissues including muscle and colon. If included, it also induces a PTC downstream. It is surrounded by a PCCR formed by R3 and R4 with Δ*G*=-34.3 kcal/mol. Exon 5b is located in between bromodomains 1 and 2.

### Transcript structure and poison exons

The human *BRD2* gene spans 13 exons, with the start codon located in exon 2. The inclusion of the 92-nt-long exon 3b, which follows exon 3, causes a frameshift that generates a premature termination codon (PTC) in the downstream exon (Figure 1B). Our bioinformatic analysis indicated that exon 3b is located inside a PCCR (id 668329) formed by regions R1 and R2 with the hybridization free energy Δ*G* = −29.1 kcal/mol [26]. Although the level of nucleotide sequence variation is insufficient to estimate the significance of compensatory substitutions in R1 and R2, their boundaries strongly correlate with the phylo-HMM profile forming two pronounced phastCons peaks, which disappear when the complementarity ends — a pattern that has been reported previously for many functional RNA structures [27].

This observation naturally led us to ask whether *BRD3*, the closest paralog of *BRD2*, also contains a poison exon. In examining the homologous part of *BRD3*, no poison exon was found in between exons 3 and 4, however we identified a conserved element in the intron spanning between exons 5 and 6 (Figure 1C). This element represents a 82-nt-long unannotated exon 5b with the canonical GT/AG splice site consensus sequences, which is expressed at low levels in GTEx tissues, namely in muscle and colon (see Methods). Inclusion of this exon also results in a frameshift that generates a PTC downstream. Furthermore, this cryptic exon is surrounded by a pair of standalone phastCons peaks (regions R3 and R4) strongly resembling those observed in *BRD2*. Upon inspection of the nucleotide sequences of R3 and R4, we found that they are complementary with Δ*G* = −34.3 kcal/mol and have a characteristic, abrupt decrease of phylo-HMM score beyond the base-paired region. Interestingly, this PCCR was not in scope of earlier studies because it fell outside the conserved RNA elements track [26, 28].

The multiple alignment of protein aminoacid sequences of the major protein-coding human BET protein isoforms reveals a remarkable conservation of exon boundaries (Figure S3). However, exon 3b of *BRD2* and exon 5b of *BRD3* reside in non-orthologous introns, one separating exons encoding a bromodomain and the other located in between bromodomains (Figure 2). We examined the *BRD4* transcript and found an annotated poison exon 3b, however pertaining to yet another intron with respect to those containing poison exons in *BRD2* and *BRD3*. This poison exon lacks any conserved RNA structure in the flanking introns (Figure S4). The last and the most distant member of the BET family, *BRDT*, neither has annotated NMD isoforms, nor contains any expressed conserved intronic element that could possibly represent a poison exon. Interestingly, the exon 3b of *BRD2* can be traced back to amphibians, while poison exons in *BRD3* and *BRD4* first appear in mammals (Figure S5).

**Figure 2:**
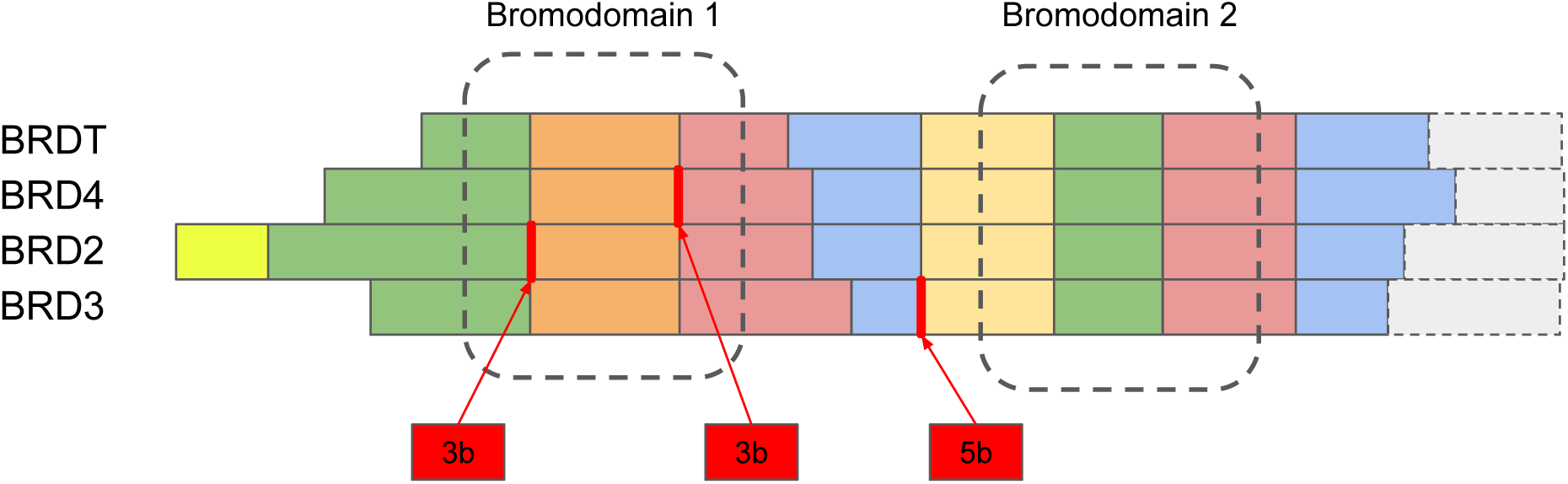
Conservation of exon-intron structure in the BET family. The schema represents a multiple sequence alignment of the human BET family members (full alignment in Figure S3). Colored blocks represent different exons. Gray rectangles denote the rest of the protein. Regions encoding bromodomains 1 and 2 are shown by dashed lines. Red vertical lines represent splice junctions corresponding to poison exons (the latter are shown as red rectangles below).

In sum, we identified two stable PCCRs surrounding poison exons in *BRD2* and *BRD3*, of which the latter exon has not been annotated before. These poison exons reside in introns spanning between non-homologous protein-coding exons, which suggests their independent acquisition in the course of evolution.

### RNA structure and poison exon inclusion

To test the effect of RNA structure on the inclusion of poison exons in *BRD2* and *BRD3*, we followed the strategy outlined previously [29, 30]. Namely, we measure splicing changes in the endogenous pre-mRNA in response to the blockage of PCCRs by antisense oligonucleotides (AONs), and construct minigenes carrying mutated gene fragments to assess splicing outcomes in single and double mutants.

First, we designed an AON complementary to the R1 sequence in *BRD2* (Figure 3A) and measured the rate of exon inclusion upon dose-dependent AON treatment by RT-PCR and qPCR. Exon 3b inclusion rate significantly increased even at the low 25-nM concentration of AON1 (Figures 3B and S6A). Next, we assembled minigene constructs carrying a fragment of *BRD2* between exons 3 and 4 (Figure 3C) with single disruptive mutations and a double compensatory mutation (Figure 3D). Transfection of these constructs into A549 cells revealed that disruptive mutations (m1 and m2) significantly increased exon 3b inclusion, while the double compensatory mutation (m1m2) reversed splicing back to that of the wild type (Figures 3E and F).

**Figure 3:**
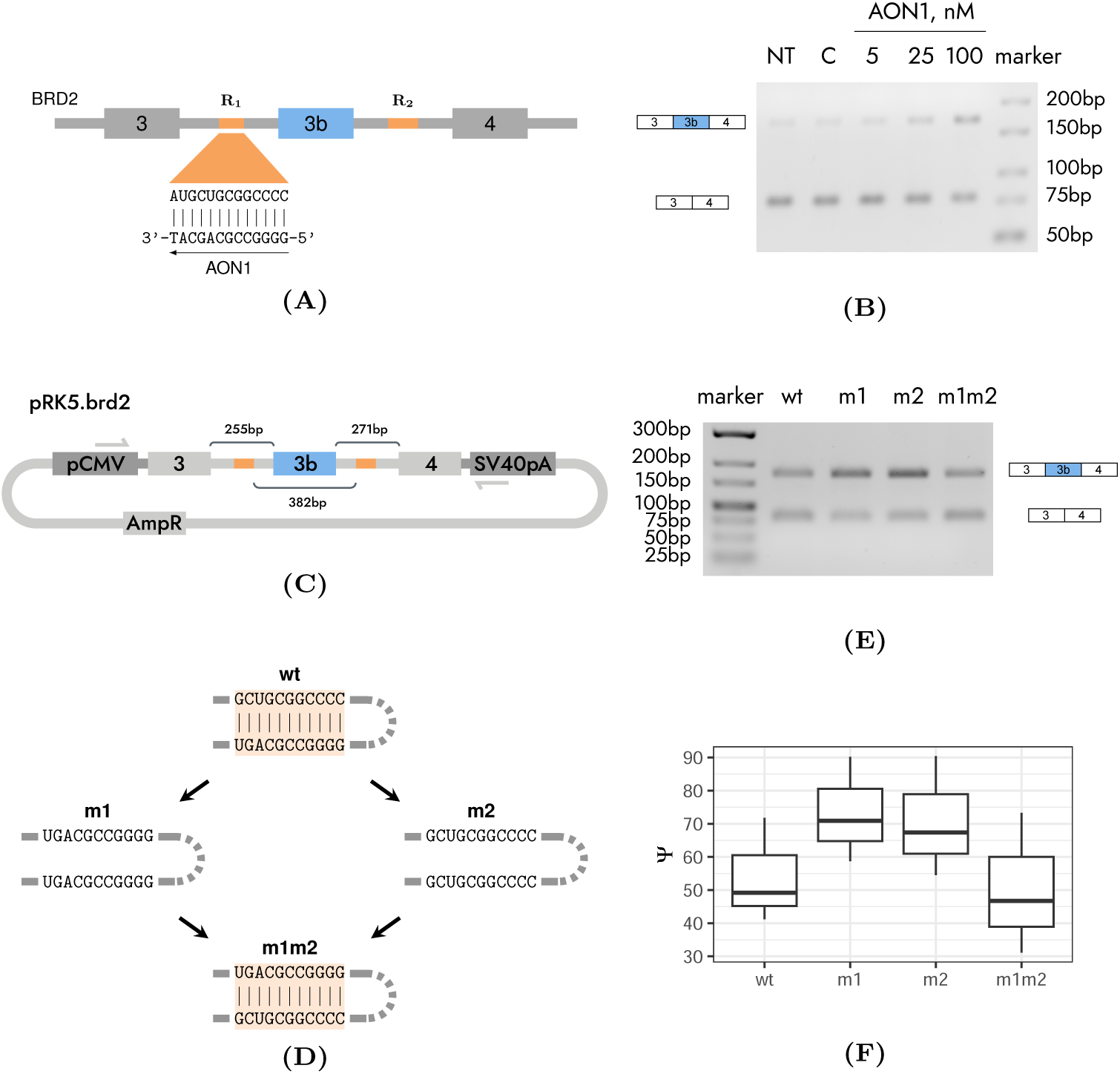
Validation of the RNA structure in the *BRD2* gene. **(A)** AON1 is complementary to the region R1. **(B)** Exon 3b inclusion rate increases when R1/R2 base pairings are blocked by AON1. ‘NT’ denotes non-treated control, ‘con’ denotes the treatment with control AON against luciferase. AON concentrations are in nM. **(C)** The minigene pRK5.BRD2 carrying a fragment of *BRD2* between exons 3 and 4. pCMV promoter and SV40 early polyadenylation signal are shown. **(D)** Mutagenesis strategy is to create single disruptive mutants (m1 and m2) and a double compensatory mutant (m1m2). **(E, F)** Exon 3b inclusion rate significantly increases in disruptive mutants but reverts to the wild type (WT) in the compensatory double mutant. A representative gel electrophoresis is shown).

In *BRD3*, the treatment with AON complementary to the R3 sequence (Figure 4A) induced higher exon 5b inclusion rate as evidenced by RT-PCR (Figure 4B) and qPCR (Figure S6B).

**Figure 4:**
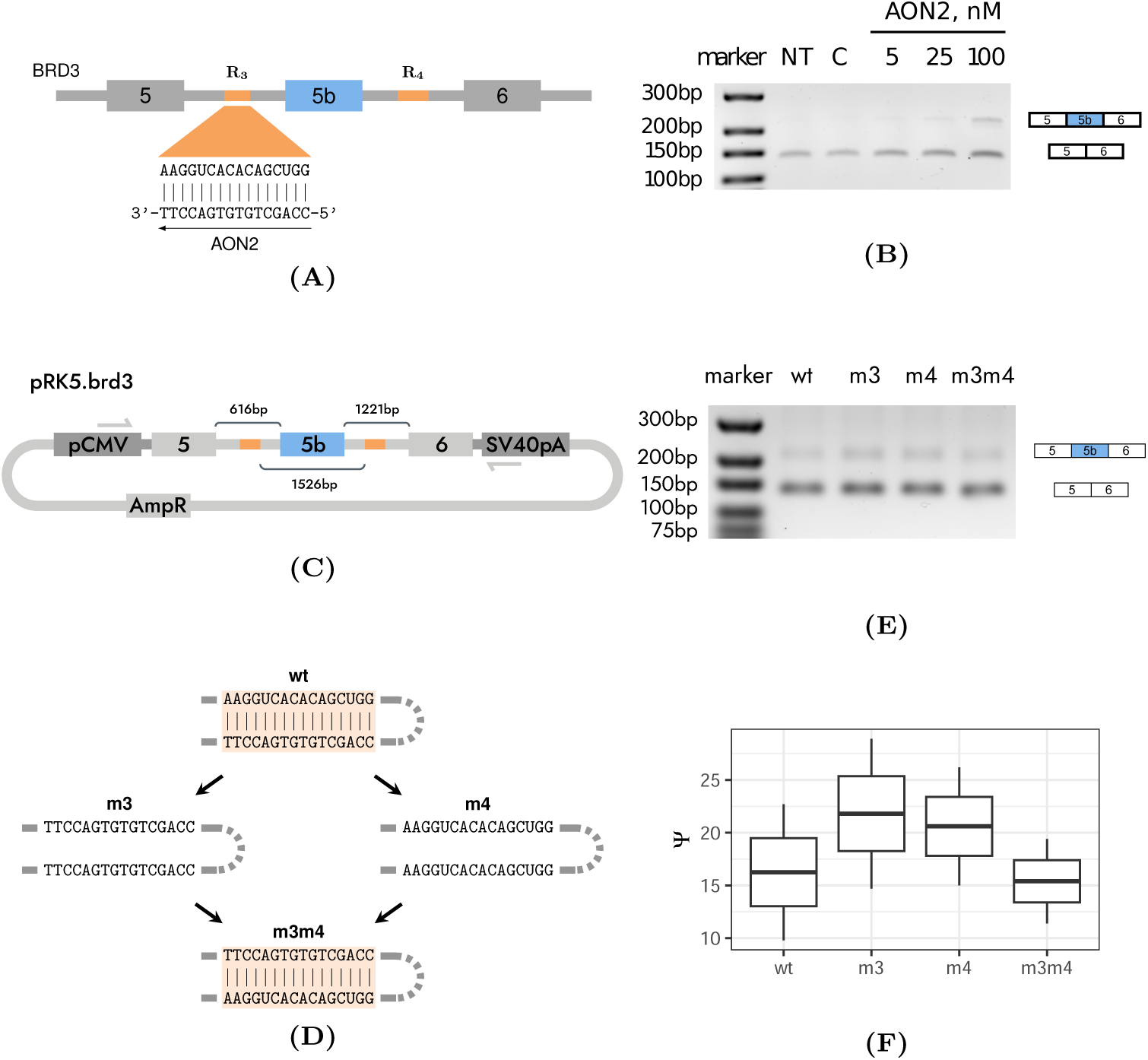
Validation of the RNA structure in *BRD3* gene. **(A)** AON2 is complementary to the region R3. **(B)** Exon 5b inclusion rate increases when R3/R4 base pairings are blocked by AON2. **(C)** The minigene pRK5.BRD3 carrying a fragment of *BRD3* between exons 5 and 6. **(D)** Mutagenesis strategy is to create single disruptive mutants (m3 and m4) and a double compensatory mutant (m3m4). **(E, F)** Exon 5b inclusion rate significantly increases in disruptive mutants but reverts to the wild type (WT) in the compensatory double mutant. The notation in the legend is as in Figure 3.

Transfection of A549 cells with minigene constructs carrying the respective fragment of *BRD3* (Figure 4C) with disruptive (m3 and m4) and compensatory (m3m4) mutations (Figure 4D) again indicated that exon 5b inclusion rate increases when base pairings are disrupted, but this increase is reversed in the compensatory double mutant (Figures 4E and F). The cryptic origin of exon 5b in *BRD3* is reflected in the observation that transcript isoforms containing this exon have low expression levels. Taken together, these experiments demonstrate that base pairings within R1/R2 and R3/R4 control the rate of inclusion of *BRD2* exon 3b and *BRD3* exon 5b, respectively.

### Expression and splicing of poison exons

Panels of large-scale transcriptomic experiments offer a powerful tool for exploring gene expression and splicing profiles across clinically relevant conditions. Accordingly, we first characterized the patterns of BET genes’ expression and splicing in healthy human tissues (GTEx). The log_10_-transformed median abundance of *BRD2* transcripts (log_10_ *TPM*) in a tissue was negatively correlated with the median level of exon 3b inclusion (Ψ) in that tissue (*r_p_* = −0.34, Figures 5A), and the negative trend was evident despite variation across samples (Figure S7). A remarkably high abundance of *BRD2* transcripts and remarkably low Ψ values were observed in testis, where human *BRD2* was reported to be consistently expressed [31].

**Figure 5:**
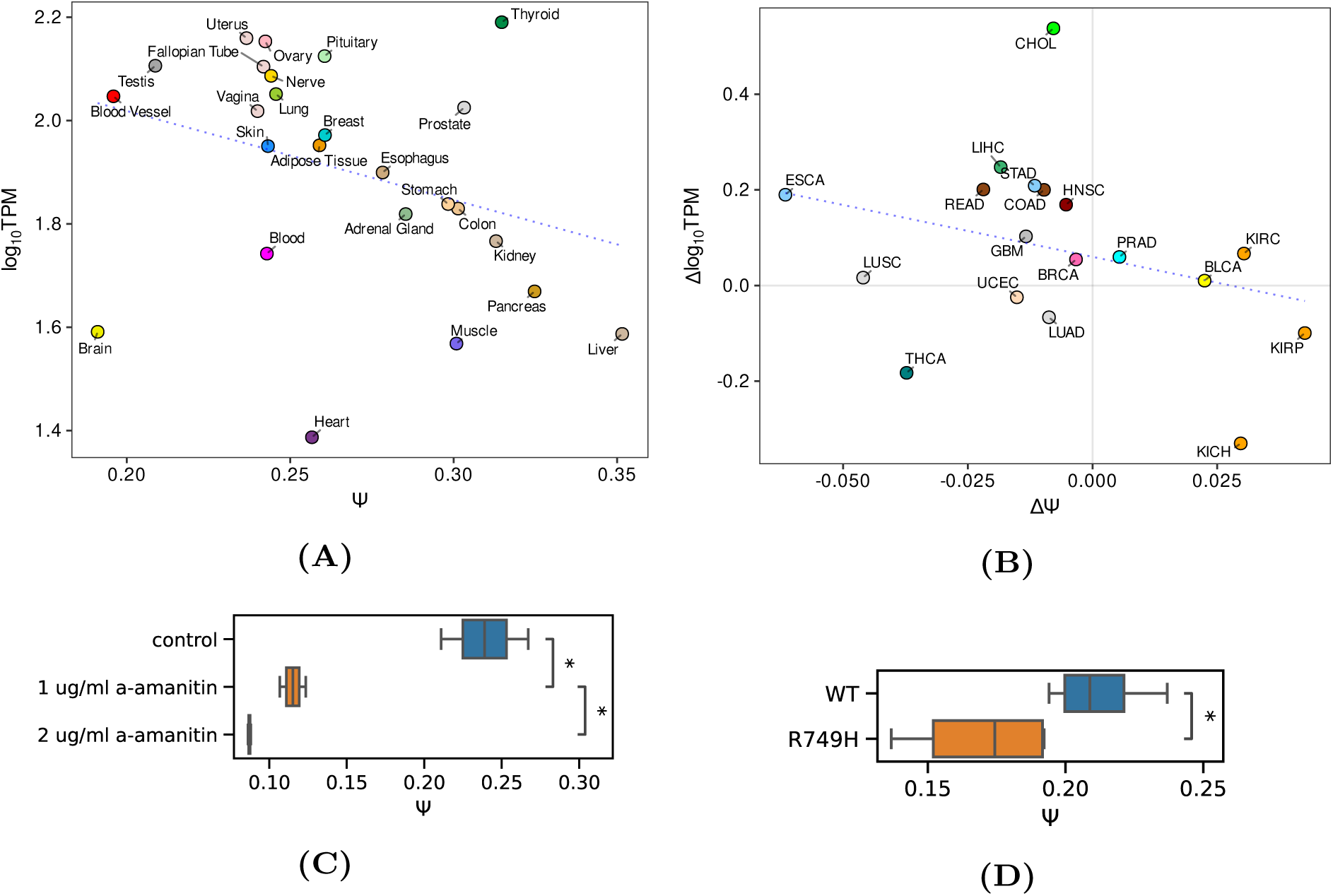
The patterns of *BRD2* expression and splicing. **(A)** The tissue-median rate of exon 3b inclusion (Ψ) is negatively associated (*r_p_* = −0.34) with the log_10_-transformed tissue-median *BRD2* transcript abundance (log_10_ *TPM*). The optimal least squares line is shown in dashed blue. Tissue color codes are as in [32]. **(B)** The difference in the median exon 3b inclusion rate (ΔΨ, tumor vs. normal tissue) is negatively correlated (*r_p_* = −0.3) with the difference in log_10_-transformed median transcript abundance (Δ log_10_ *TPM*). TCGA project color codes and abbreviations are as in [32]. **(C)** The response of exon 3b inclusion (Ψ) to α-amanitin treatment. **(D)** Exon 3b inclusion rate (Ψ) in the wild type (WT) and in the slow RNA pol II mutant (R749H).

A similar relationship was observed when comparing the *BRD2* expression and splicing levels across a transcriptomic panel of human cancers (TCGA). The difference in log_10_-transformed median transcript abundance (Δ log_10_ *TPM*, tumor vs. normal tissue) was negatively correlated (*r_p_* = −0.3) with the respective difference in the median exon 3b inclusion rate (ΔΨ) across eighteen cancer types analyzed (Figures 5B and S8). Remarkably, the highest increase in exon 3b inclusion accompanied by a substantial drop in *BRD2* expression were observed in clear cell carcinoma of the kidney (KIRC) and renal papillary cell carcinoma (KIRP), two cancer types, in which the downregulation of *BRD2* was shown to be associated with poor survival rate [33].

While these observations are in agreement with the poison role of exon 3b, which should expectedly lead to a decrease in transcript abundance with its inclusion, a question remains whether RNA structure R1/R2 is involved in the regulation of exon 3b splicing. We addressed this question by examining two available datasets on the transcriptome response to the transcription elongation slowdown, one under α-amanitin treatment [26] and the other in the slow RNA Pol II mutant R749H [34]. In both cases, the inclusion of exon 3b significantly declined (P-value = 0.002 and P-value = 0.02, respectively) when RNA Pol II speed was slowed down (Figures 5C and D). In accordance with poison exon skipping, the expression level of *BRD2* expectedly increases, however the difference was not statistically significant (P-value = 0.15).

Poison exons in *BRD3* and *BRD4* are expressed at much lower levels as compared to *BRD2* exon 3b (Figure S9), and their assessment is possible only in a fraction of tissues. In *BRD3*, the median transcript abundance and the median exon 5b inclusion rate are negatively associated (*r_p_* = −0.25), while in *BRD4* the relationship is positive (*r_p_* = 0.22), but driven mostly by an influential score representing testis tissue (Figure S10). The response of poison exons in *BRD3* and *BRD4* to RNA Pol II slowdown cannot be reliably estimated due to low short read coverage.

## Discussion

In recent years, experimental and computational studies have demonstrated that RNA structure is critically important for alternative splicing regulation [35, 29, 26]. While the overall function of alternative splicing can arguably be attributed to expanding the proteome diversity [36, 37, 38], the functional implication of altering the primary sequence of a particular protein in most cases remains undocumented, thereby limiting the interpretation of RNA structure’s impact. Here, for the first time, we identified two PCCRs that control unproductive splicing representing an important example, in which the function of RNA structure can be directly attributed to posttranscriptional regulation of gene expression rather than to changes in the protein product itself.

Paralog genes that arise through duplication and divergence often maintain a considerable degree of similarity not only between their encoded proteins but also between cis-regulatory elements and regulatory control mechanisms [39]. In particular, this is true for serine-arginine-rich (SR) splicing factors, many of which evolved through duplications and contain homologous cis-regulatory elements that are associated with unproductive splicing [19]. On the other hand, partial duplication of complementary sequences represents a general evolutionary mechanism for the generation of competing RNA structures that regulate mutually exclusive splicing [40]. Therefore, genomic duplications naturally occur at the crossroads of unproductive splicing and RNA structure evolution.

Surprisingly, in the case of *BRD2* and *BRD3*, the regulatory poison exons are located in introns spanning between non-homologous exons. This suggests that *BRD2* and *BRD3* either acquired them independently, or each independently lost one of them in the evolution from the common ancestor. The fact that *BRD4* contains a poison exon but doesn’t have RNA structure around it, and furthermore that *BRDT* and *fs(1)h*, an invertebrate homolog of BET proteins, don’t have any annotated NMD isoforms strongly indicated that the former scenario took place. Furthermore, the conserved complementary sequences surrounding poison exons in *BRD2* and *BRD3* share no sequence similarity with each other. All these observations suggest that the trait represented by regulatory RNA structures around poison exons is a result of convergent evolution. Rapid loss and reacquisition of poison exons [23] and independent origins of competing RNA secondary structures that control mutually exclusive splicing of tandemly duplicated exons [41] suggest that convergent evolution of RNA structures regulating unproductive splicing may extend far beyond the examples presented here.

In searching for the driving force of this convergent evolution, one may conjecture that RNA structures could regulate unproductive splicing through a mechanism related to transcription elongation rate. Slower RNA Pol II elongation rate allows sufficient time for RNA structure to fold, which, in turn, could lead to skipping of the looped-out exon. Such a mechanism was reported for the *ATE1* gene, which contains a pair of ultra-long-range complementary regions that control splicing of mutually exclusive exons [30]. NELFE, a factor linked to RNA Pol II pausing such as [42, 43, 44], could be responsible for poison exon skipping and increased expression of *BRD2* and *BRD3* in testis. SR-rich splicing factors often implement a negative feedback loop that downregulates the productive splice-isoform in response to elevated expression of the gene product [45]. A similar feedback loop may also exist in *BRD2*, however acting through alternative splicing rather than RNA Pol II slowdown since *BRD2* does not appear to alter the transcription elongation rate itself [46].

The example of *BRD2* demonstrates that regulatory RNA structure may extend beyond PCCRs reported here, as the intron downstream of exon 3b contains a polymorphic microsatellite, the number of GT-repeats in which negatively correlates with the rate of exon 3b inclusion [21]. In general, self-base-pairing sequences such as GT tracts strongly affect splicing rate due to RNA secondary structure formation [47]. While RNA structures formed by R1/R2 and R3/R4 were identified on the basis of evolutionary conservation, base pairings that are important for unproductive splicing may well exist in mRNAs of other BET family members, however outside of conserved regions. This, and the extent to which RNA structure fits into the complex regulatory landscape of unproductive splicing and long ultraconserved elements associated with it, remains a question for future studies.

Studying mechanisms underlying BET-specific functions have far-reaching therapeutic implications as abnormal expression of these proteins leads to oncological, metabolic, and cardio-vascular disorders [48, 49, 50]. The approach based on synthetic AONs was proven effective in targeting unproductive splicing, particularly in bromodomain proteins. For instance, AONs that promote skipping of poison exon 14a, which lead to increased *BRD9* mRNA degradation in tumors, help to restore *BRD9* protein levels and reverse tumor growth [22]. In contrast, AONs targeting RNA structure elements in *BRD2* and in *BRD3* offer a unique opportunity to not only suppress but also increase poison exon inclusion therapeutically. More importantly, the evidence for convergent evolution of poison exons and RNA structures in *BRD2* and in *BRD3* provides an orthogonal view on the mechanisms of unproductive splicing regulation in these and many other paralogous genes.

## Conclusion

This study for the first time describes RNA structures that regulate unproductive splicing. Poison exons in *BRD2* and *BRD3*, of which the latter is a cryptic exon and wasn’t annotated before, represent an example of convergent evolution that led to independent acquisition of poison exons and regulatory RNA structures. This study provides important implications for selective modulation of *BRD2* and *BRD3* expression in therapeutic applications.

## Methods

### RNA-seq data

RNA-seq data of poly(A)^+^ RNA obtained in the Genotype-Tissue Expression project (GTEx) were downloaded from dbGaP (dbGaP project 15872) in fastq format and aligned to the human genome assembly GRCh38 (hg38) using STAR aligner version 2.7.3a in paired-end mode [51] with GENCODE v42 annotation as a reference [52]. The RNA-seq data in RNA Pol II slowdown experiment with α-amanitin and in RNA pol II mutants with slow elongation rate were downloaded from GEO repositories GSE153303 and GSE63375, respectively [26, 34]. The gene read coverage was calculated from BAM files using the featureCounts tool from Subread package version 2.0.6 [53] in paired-end mode (-p --countReadPairs options). TPM (transcripts per million) values were calculated from the read coverage using RNAnorm package version 2.0.0. Split read counts and percent-spliced-in metrics were computed by IPSA pipeline [54] with default settings as before [29]. Namely, the exon inclusion rate (Ψ) was computed as, 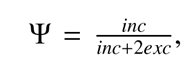 where *inc* is the number of split reads supporting exon inclusion, and *exc* is the number of split reads supporting exon exclusion. Ψ values with the denominator below 10 were discarded.

### Cryptic exons

To search for cryptic exons, BAM files from GTEx were processed with StringTie [55] in guided assembly mode with conservative parameters (--conservative option) using GENCODE v42 annotation. Two groups of non-terminal exons were selected from the StringTie output: (1) annotated exons and (2) exons that don’t intersect any annotated exon from GENCODE. The following metrics were computed for each exon in each sample: the read coverage per base pair, the total number of split reads supporting splice junctions connecting the exon with any annotated exon, and the average PhastCons score derived from the multiple alignment of 100 vertebrate genomes [56]. The 15^th^ percentiles of these metrics for group (1) were used as a cutoff for group (2). This procedure yielded 249 cryptic (unannotated) exons with all three metrics exceeding the cutoffs chosen for group (1).

### Phylogenetic analysis

To construct the phylogenetic tree, we selected the smallest orthologous group in the eggNOG database including the human proteins *BRD2*, *BRD3*, *BRD4*, and *BRDT* (ENOG503CSGG for the taxon Bilateria) [24]. All nodes corresponding to homologs from *H. sapiens* and *D. Rerio* were selected. The *fs(1)h* protein from *D. Melanogaster* was used as an outgroup. The pruned tree was visualized with IToL web service [57]. The aminoacid sequences of the major proteincoding isoforms of the human BET proteins were obtained from the Ensembl database [58] and aligned using Muscle software v3.8.1551 [59]. The alignment and the phylogenetic trees constructed for bromodomains 1 and 2 were visualized using Jalview editor [60].

### Statistical procedures

The data were analyzed using python version 3.8.2 and R statistics software version 3.6.3. Throughout the paper, *r_p_* denotes the Pearson correlation coefficient. The significance of *r_p_* values was not assessed due to a small number of observations (tissues and TCGA projects). Non-parametric tests were performed using normal approximation with continuity correction. In all figures, the significance levels 0.05, 0.01, and 0.001 are denoted by *, **, and ***, respectively.

### Cell culture and transfection

Human A549 lung adenocarcinoma cells were maintained in Dulbecco’s modified Eagle’s medium Nutrient Mixture F-12 with phenol red (Thermo Fisher Scientific or Servicebio) supplemented with 10% v/v fetal bovine serum (FBS) and 1% GlutaMAX (Thermo Fisher Scientific). All treatments were done in a 12-well plate format. A549 cells were plated at a density of 150,000 cells per well. Five hundred nanograms of WT or mutated minigene plasmids were transfected into A549 cells using Lipofectamine 3000 (Invitrogen) for 24 hours. AON (13-mer) transfection was performed with Lipofectamine RNAiMAX (Invitrogen) in OptiMEM serum-reduced media (Gibco) for 48 hours. Each experiment was made in at least three independent biological replicates.

### Minigene construction and mutagenesis

A fragment of the human genome from exon 5 to 6 of the *BRD3* gene was amplified using Q5 High-Fidelity DNA Polymerase (New England Biolab). The genomic DNA of the human A549 lung adenocarcinoma cell line was used as a template. All sequences were inserted downstream of the cytomegalovirus (CMV) promoter of the pRK5 vector. The minigene encoding *BRD3* exons 5–6 was made using the NEBuilder HiFi DNA Assembly Master Mix (New England Biolab). The minigene encoding *BRD2* exons 3–4 was constructed using the blunt-end cloning approach. Mutations in all minigenes were introduced by PCR-based site-directed mutagenesis. Primers for cloning and mutagenesis are listed in Tables S1 and Tables S2, respectively. All constructs were confirmed by sequencing and restriction analysis.

### RNA extraction

Total RNA was isolated by a guanidinium thiocyanate phenol-chloroform method using ExtractRNA Reagent (Evrogene) or PureLink RNA minikit (Invitrogen) [61]. One thousand ng of total RNA was first subjected to RNase-Free DNase I digestion (Thermo Fisher Scientific) at 37°C for 30 minutes to remove contaminating gDNA. Next, 500 ng of total RNA was used for cDNA synthesis using Maxima First Strand cDNA Synthesis Kit for RT-qPCR (Thermo Fisher Scientific) to a final volume of 10 µl. cDNA was diluted 1:10 with nuclease free water for qPCR and RT-PCR analysis.

### RT-PCR and RT-qPCR

RT-PCR analysis was used to assess the ratio of splice isoforms as described elsewhere [29]. qPCR reactions were performed in triplicates in a final volume of 12µl in 96-well plates with 420 nM gene-specific primers, 2µl of cDNA using 5X qPCRmix-HS SYBR reaction mix (Evrogen). Primers for RT-PCR and RT-qPCR are listed in Tables S3. A sample without reverse transcriptase enzyme was included as control to verify the absence of genomic DNA contamination. Amplification of the targets was carried out on CFX96 Real-Time system (Bio-Rad), with following parameters: 95°C for 2 min, followed by 45 cycles at 95°C for 30 s, 61°C for 30 s, and 72°C for 30 s, ending at 72°C for 5 min. Gene and gene isoform expression change was calculated using an estimate of the amplification efficiency value. To account for the relation between pipetting error and lower copy number of template input in the reaction we discarded all the replicates that don’t reside within acceptable *C_q_*range as described in [62].

### Antisense oligonucleotides

All antisense oligonucleotides (AONs) were designed as locked nucleic acid (LNA)-based with a DNA substitution at every second nucleotide [63]. Synthesis of LNA/DNA mixmers was carried out by Syntol JSC (Moscow, Russia). Sequences of AONs are listed in Table S4.

## Authors’ contributions

DP designed and supervised the study; DP and SM performed data analysis; MP, MV, and DS performed the experiments; DP, SM, and MP wrote the first draft of the manuscript. All authors edited the final version of the manuscript.

## Acknowledgments

The results presented here are in part based on the data generated by the TCGA Research Network (https://www.cancer.gov/tcga) and data obtained from the GTEx Portal and dbGaP accession number phs000424/GRU on 12/10/2018. The authors acknowledge Prof. Sergei Spirin for insightful discussions on BET family phylogeny.

