## Supplementary Information for "BRD2 and BRD3 genes independently evolved RNA structures to control unproductive splicing"

October 7, 2023

### List of Figures

### List of Tables

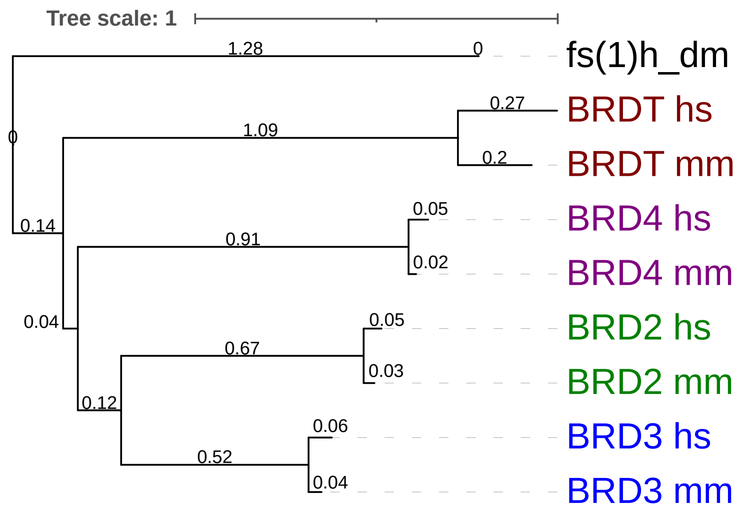


Figure S1: The phylogenetic tree of bilaterian homologs of *BRD2* from eggNOG database restricted to *H. Sapiens* and *M. musculus*. The fs(1)h protein from *D. Melanogaster* was used as an outgroup. The numbers on the branches denote the average number of aminoacid substitutions per site.

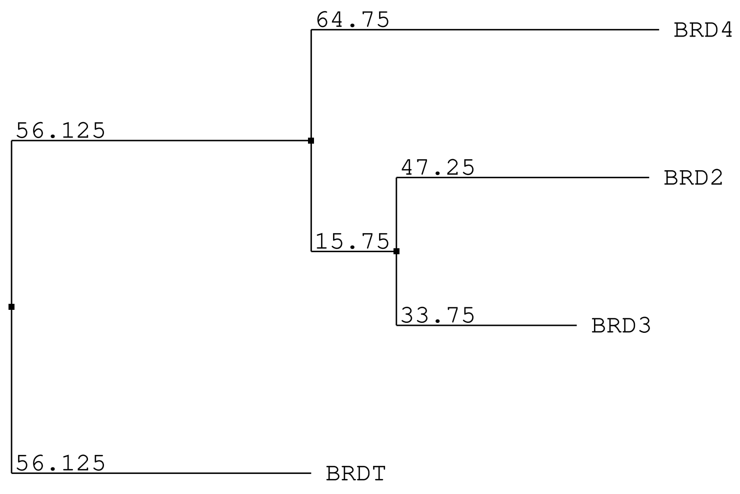


(A)

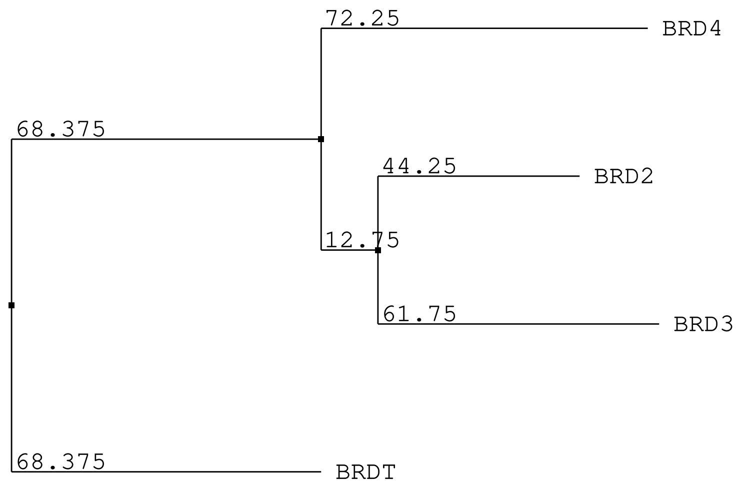


(B)

Figure S2: Phylogenetic trees of the human *BRD2*, *BRD3*, *BRD4*, and *BRDT* constructed from the alignment of bromodomain 1 (A) and bromodomain 2 (B). The numbers on the branches represent bootstrap values.

```
BTDT/1-947       -----------------------------------------------------mslpsrq
BRD4/1-1362      ----------------------msaesgpgtrlrnlpvmgdgletsqmsttqaqaqpqpa
BRD2/1-801       ------mlqnvtphnKlpgegnagllglgpeaaapgkrirkpsllyegfesptmasvpal
BRD3/1-726       ----------------------------------------------mstattvapagipa

                                  |<=========================================
BTDT/1-947       taiivnppppeyintkkngrltnqlqylqkvvlkdlwkhsfswpfqrpvdavklqlPDyy
BRD4/1-1362      naastnppppetsnpnkpkrqtnqlqyllrvvlktlwkhqfawpfqqpvdavklnlPDyy
BRD2/1-801       qltpanppppevsnpkkpgrvtnqlqylhkvvmkalwkhqfawpfrqpvdavklglPDyh *
BRD3/1-726       tpgpvnppppevsnpskpgrktnqlqymqnvvvktlwkhqfawpfyqpvdaiklnlPDyh

                 ================      BROMODOMAIN1     =====================
BTDT/1-947       tiiknpmdlntikkrlenkyyakaseciedfntmfsncylynKPgddivlmaqaleklfm
BRD4/1-1362      kiiktpmdmgtikkrlennyywnaqeciqdfntmftncyiynKPgddivlmaealeklfl *
BRD2/1-801       kiikqpmdmgtikrrlennyywaasecmqdfntmftncyiynKPtddivlmaqtlekifl
BRD3/1-726       kiiknpmdmgtikkrlennyywsasecmqdfntmftncyiynKPtddivlmaqalekifl

                 =======>|
BTDT/1-947       qklsqmpqeeq---vvgvkerikk----Gt-------qqniavssakeksspsatek---
BRD4/1-1362      qkinelpteeteimivqakgrgrgrketGtakpgvstvpnttqastppqtqtpqpnpppv
BRD2/1-801       qkvasmpqeeqelvvtipknshkkgaklaAlqgsvtsahqvpavssvshtalytp-ppei
BRD3/1-726       qkvaqmpqeevellppapkgkgrkpaagaqsaGt----qqvaavssvspatpfqsvpptv

BTDT/1-947       -------------vfkqqeipsvfpk------tsisplnvvqgasvnsssqtaa------
BRD4/1-1362      qatphpfpavtpdlivqtpvmtvvppqplqtpppvppqpqpppapapqpvqshppiiaat
BRD2/1-801       ptt-----------vlniphpsviss-------pllkslhsagppll-avtaapp----a
BRD3/1-726       sqtp---------viaatpvptitan---vtsvpvppaaappppatp-ivpvvpp----t

BTDT/1-947       ----QVtkgvkrkadtttpat-savk-assefsptf-teksvalppikenm-----pknv
BRD4/1-1362      pqpvKTkkgvkrkadtttpttidpih-eppslpp--epkttklgqrress-rpvkppkkd
BRD2/1-801       qplaKKkgvkrkadtttptptailapgspasppgslepkaarlppmrresgrpikpprkd
BRD3/1-726       ppvvKKkgvkrkadtttpttsaita-srsespppplsdpkqakvvarresggrpikppkkd *

                                |<==========================================
BTDT/1-947       lpdsqqqynvvktvkvteqlrhcseilkemlakkhfsyawpfynpvdvnalglhnyydvv
BRD4/1-1362      vpdsqqhpapeksskvseqlkccsgilkemfakkhaayawpfykpvdvealglhdycdii
BRD2/1-801       lpdsqqqhqsskkgklseqlkhcngilkellskkhaayawpfykpvdasalglhdyhdii
BRD3/1-726       ledgevpqhagkkgklsehlrycdsilremlskkhaayawpfykpvdaealelhdyhdii

                 ================      BROMODOMAIN2     =====================
BTDT/1-947       knpmdlgtiKEkmdnqeykdaykfaadvrlmfmncykynppdhevvtmarmlQDvfethf
BRD4/1-1362      khpmdmstiKSkleareyrdaqefgadvrlmfsncykynppdhevvamarklQDvfemrf
BRD2/1-801       khpmdlstvKRkmenrdyrdaqefaadvrlmfsncykynppdhdvvamarklQDvfefry
BRD3/1-726       khpmdlstvKRkmdgreypdaqgfaadvrlmfsncykynppdhevvamarklQDvfemrf

                 ===>|
BTDT/1-947       skipiepvesmpl---cyiktditettgrentneass-----------------egnssd
BRD4/1-1362      akmpdep-eepvvavsspavppptkvvappsssdsssdsssd------------sdsstd
BRD2/1-801       akmpdeplepgplpvstamppglaksssessseessesssseeeeeedeedeeeeesess
BRD3/1-726       akmpdepveapalp--apaapmvskgaessrsseess-----------------sdsgss

BTDT/1-947       dsedervkrlaklqeQLkavhqqlqvlsqvpfrklnkkkekskkekk---------kekv
BRD4/1-1362      dseeeraqrlaelqeQLkavheqlaalsqpqqnkpkkkekdkkekkk----ekhkrkeev
BRD2/1-801       dseeerahrlaelqeQLravheqlaalsqgpiskpkrkrekkekkkkrka-ekhrgraga
BRD3/1-726       dseeeratrlaelqeQLkavheqlaalsqapvnkpkkkkekkekekkkkdkekekekhkv

BTDT/1-947       nnsnenprkmceqmrlkek----------------------skrNqpkkrkqqf-----i
BRD4/1-1362      eenkkskakepppkktkknns--------------------snsnvSkkepapmkskppp
BRD2/1-801       deddkgpraprppqpkkskkasgsgggsaalgpsgfgpsggsgtKlpkkatktappalpt
BRD3/1-726       kaeeekkakvappakqaqqkkapakkansttt---------agRqlkkggkqas-----a

                    |<====================== ET DOMAIN ======================
BTDT/1-947       glksedednakpmnydekrqlslninklpgdklgrvvhiiqsrepslsnsnpdeieidfe
BRD4/1-1362      tyeseeedkckpmsyeekrqlsldinklpgeklgrvvhiiqsrepslknsnpdeieidfe
BRD2/1-801       gydseeeeesrpmsydekrqlsldinklpgeklgrvvhiiqarepslrdsnpeeieidfe
BRD3/1-726       sydseeeeeglpmsydekrqlsldinrlpgeklgrvvhiiqsrepslrdsnpdeieidfe
```

Figure S3: A part of the multiple sequence alignment of human BET family member aminoacid sequences including two bromodomains and a fragment of the ET domain. The highlighted blocks represent different exons. Aminoacids adjacent to exon junctions corresponding to poison exons are marked in red.

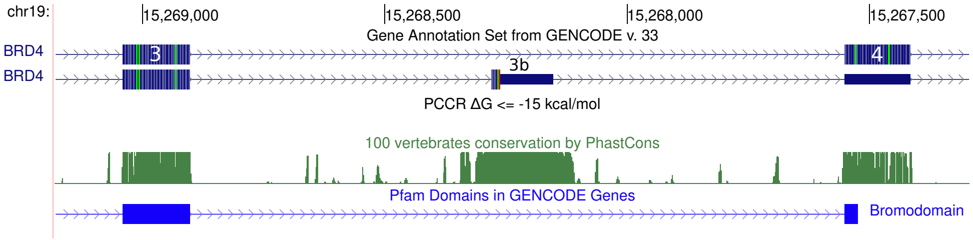


Figure S4: Poison exon in *BRD4* is not surrounded by any PCCR with $\Delta G \leq -15$ kcal/mol.

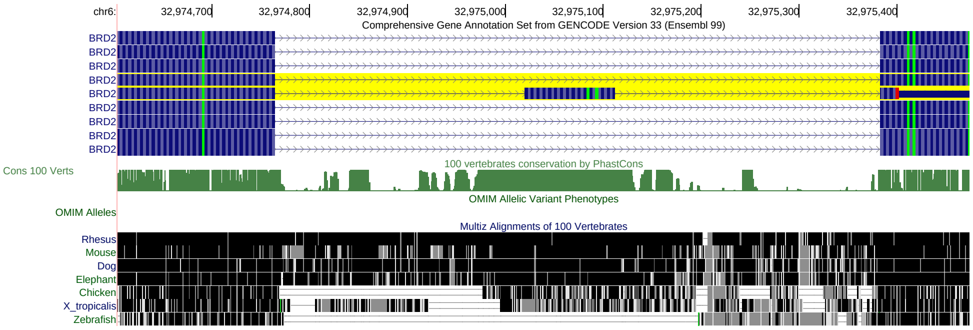


(A)

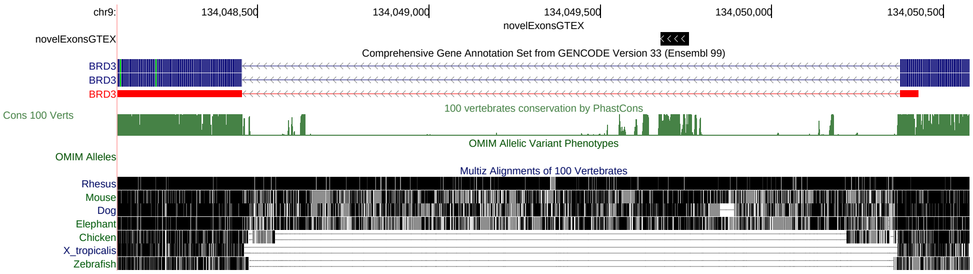


(B)

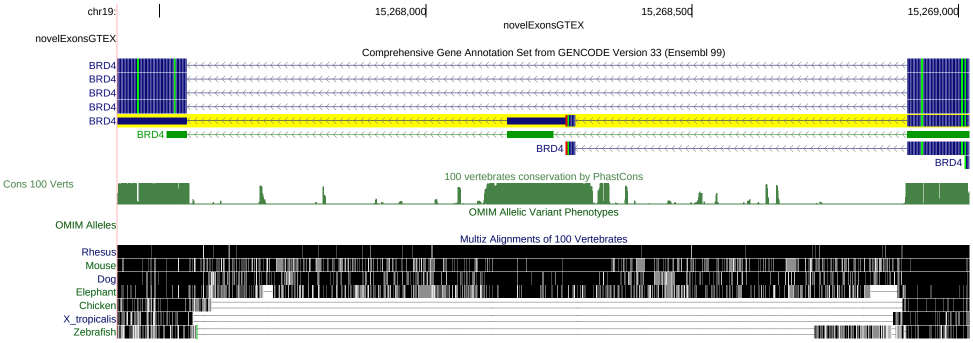


(C)

Figure S5: Poison exons in *BRD2*, *BRD3*, and *BRD4* across vertebrate evolution. In *BRD2*, poison exon 3b can be traced back to amphibians. In *BRD3* and *BRD4*, poison exons first appear in mammals.

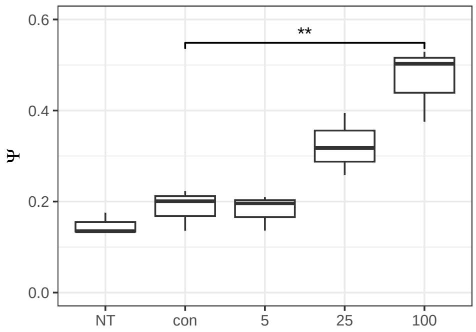


(A)

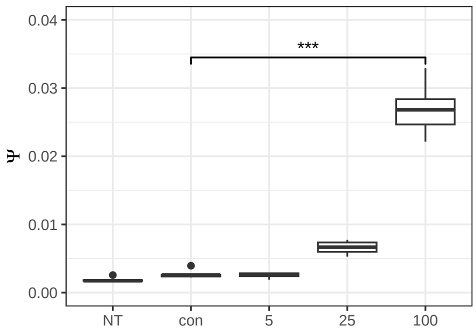


(B)

Figure S6: qPCR analysis of *BRD2* exon 3b (A) and *BRD3* exon 5b inclusion in response to AON1 (A) and AON2 (B) treatment. 'NT' denotes non-treated control, 'con' denotes the treatment with control AON against luciferase. AON concentrations are in nM. Asterisks (** and ***) denote statistically discernible differences at the 1% and 0.1% significance level, respectively.

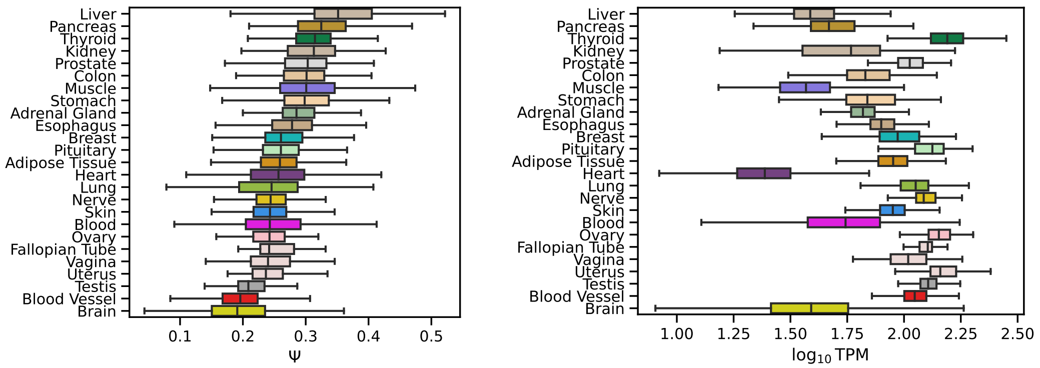


Figure S7: The distribution of Ψ, the *BRD2* exon 3b inclusion rate (left), and $\log_{10} TPM$, the *BRD2* expression level (right), across GTEx tissues.

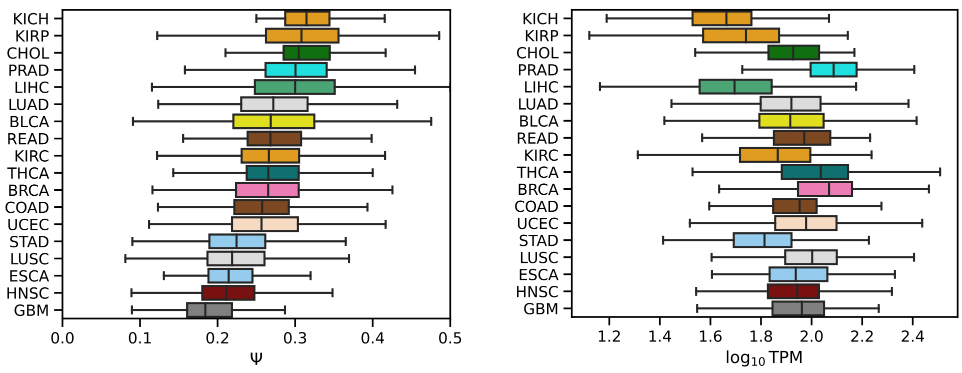


(A) (B)

Figure S8: The distribution of Ψ, the *BRD2* exon 3b inclusion rate (left), and $\log_{10} TPM$, the *BRD2* expression level (right), across TCGA cancers.

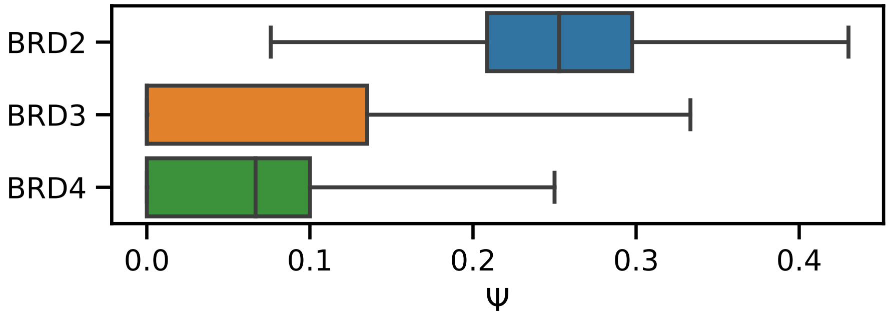


Figure S9: The inclusion rates (Ψ) of poison exons in *BRD2*, *BRD3*, and *BRD4* across GTEx tissues.

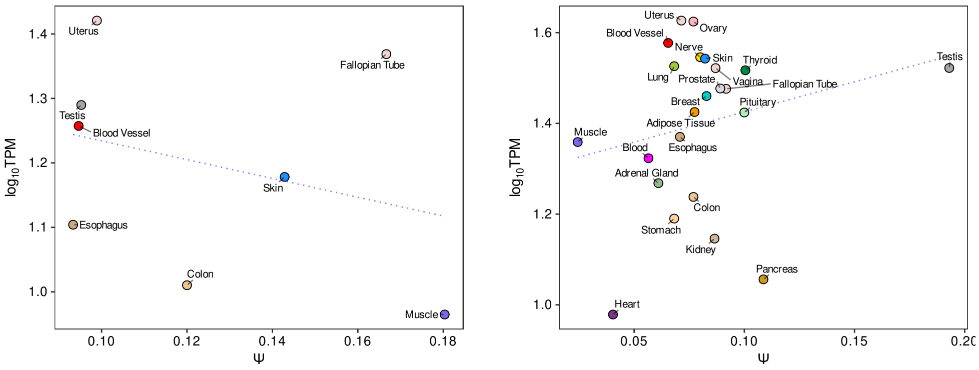


Figure S10: Left: The relationship between *BRD3* exon 5b inclusion rate (Ψ) and *BRD3* expression level ($\log_{10} TPM$). Right: The relationship between *BRD4* exon 3b inclusion rate (Ψ) and *BRD4* expression ($\log_{10} TPM$).

| | |
|---|---|
| brd2_cl_fw | CAAGGTAGTGATGAAGGCTCTG |
| brd2_cl_rv | CATACACTCTGAAGCAGCCC |
| brd3_nb_fw | ttctatcgattgaattccccGTACACAGCAAGTGGCGG |
| brd3_nb_rv | gtcgactctagaggatccccTTTCACGGTGCTGAGGTC |
| pRK5_fwd | GGGGATCCTCTAGAGTCGACCTGC |
| pRK5_rev | GGGGAATTCAATCGATAGAACCGAGG |

Table S1: Cloning primers.

| | |
|---|---|
| brd2_m1_fw | CGACGCCGGGGTAGGGAGTTCCCATTTCTC |
| brd2_m1_rv | ATCTACACTAGGCAGACCACC |
| brd2_m2_fw | CCCCGGCGTCATTAAGTAACTGTTCCTTTGATGC |
| brd2_m2_rv | AATTAAACTGTGGGACAAAATAAATAAAT |
| brd3_m1_fw | TTCCAGTGTGTCGACCTAAGGGAGGGCCTGGCTGAAC |
| brd3_m1_rv | CAGCTCCCCAGGCCTAGGT |
| brd3_m2_fw | GGTCGACACACTGGAAGGGCCAGTGACATGGCCC |
| brd3_m2_rv | CAGTGGTGAGGCTGGGACTC |

Table S2: Mutagenesis primers.

| | |
|---|---|
| brd2_fw | GATGCTGTCAAACTGGGTCT |
| brd2_rv | GTCTCCTCTTAATAGTACCCATGTC |
| brd3_fw | CATCACTGCAAACGTCACGTC |
| brd3_rv | TGTCTGCTTTCCGCTTCACG |
| brd2_in | ATGAGGGAAAGGAAGAAGCTAAG |
| brd2_ex_fw | ACAAGGTAGTGATGAAGGCTC |
| brd2_ex_rv | GTGATAATCCGGTAGACCCAG |
| brd3_in | CTTGGGCGAGCCATTTAACC |
| brd3_ex | CGCCCTTTTTCTTGACGACAG |
| gapdh_fw | TGGTCTCCTCTGACTTCAACA |
| gapdh_rv | TGTTGCTGTAGCCAAATTCGT |

Table S3: RT-PCR and RT-qPCR primers.

| | |
|---|---|
| AON1 | +G*G*+G*G*+C*C*+G*C*+A*G*+C*A*+T |
| AON2 | +C*C*+A*G*+C*T*+G*T*+G*T*+G*A*+C |

Table S4: AON sequences. DNA nucleotides: G,A,T,C. LNA nucleotides (red): +G, +A, +T, +C. Phosphorothioated DNA nucleotides: G*, A*, T*, C*. Phosphorothioated LNA nucleotides (red): +G*, +A*, +T*, +C*.
